# Dynamic changes in innate immune and T cell function and composition at the nasal mucosa across the human lifespan

**DOI:** 10.1101/576744

**Authors:** Jesús Reiné, Beatriz F. Carniel, Carla Solórzano, Elena Mitsi, Sherin Pojar, Elissavet Nikolaou, Esther L. German, Angela D. Hyder-Wright, Helen Hill, Caz Hales, Lynsey Brown, Victoria Horsley, Lisa Hughes, Seher Zaidi, Victoria Connor, Ben Morton, Andrea M. Collins, Jamie Rylance, Hugh Adler, Paul S. McNamara, Daniela M. Ferreira, Simon P. Jochems

## Abstract

The very young and very old are at increased risk of serious infections, including pneumonia. This may relate to changes in the immune system as young children have limited immunological memory, while immunosenescence, inflammaging and a decreased pool of naïve immune cells are described with advanced age. How the immune system changes with age at mucosal surfaces, from where infections frequently develop, is not very clear as access to human tissue samples is limited. Therefore, we aimed to assess the composition and activation state of the immune system at the human mucosa. Here, we profiled nasal immune cells from 207 individuals between 1 to 80 years old using flow cytometry. Neutrophil and monocyte functionality were measured using whole blood assays. Levels of thirty nasal cytokines were measured from nasal lining fluid. Nasopharyngeal colonization by *Streptococcus pneumoniae* was assessed using classical microbiology and associated with immune responses. We found that young children have a striking paucity of granulocytes at the nasal mucosa compared to adults. In addition, T cell numbers at the nasal mucosa decreased progressively with age and were almost absent in older adults. While nasopharyngeal colonization by *Streptococcus pneumoniae* was associated with elevated levels of inflammation it had a limited effect on nasal immune composition, including levels of monocytes and neutrophils. These results show that the immune system at the nasal mucosal surface changes drastically with age and provides explanations for the increased susceptibility to infections in young and old age.

**Significance statement:** How the immune system changes with age is an intensive area of research, but has been primarily studied in blood. However, blood poorly reflects the immune system at the mucosa, from where infections develop. This manuscript provides a first characterization of how the composition and function of the immune system in the upper respiratory tract changes with age, providing explanations for increased susceptibility to infection in the very young and old. Furthermore, by linking mucosal and systemic measurements with pneumococcal colonization, we observed that reduced monocyte and neutrophil responses associate with the increased burden of pneumococcal colonization in children. This study highlights the need to study the immune system also at other mucosal sites in the context of aging.

## Introduction

Pneumonia is the most common infectious cause of death in children under 5 worldwide (1). Individuals with advanced age are also at progressively increasing risk of acquiring pneumonia (2, 3). Over the next two decades, the incidence of community-acquired pneumonia is expected to double in the United States as the population ages (4). How alterations in the mucosal immune system with age predispose to infections in the very young and very old remains unclear as access to samples is limited. While many studies have investigated how the immune system changes with age, most of these have been conducted in blood, which poorly reflects the immune system at mucosal surfaces (5, 6). Nonetheless, studies from blood and secondary lymphoid tissues have revealed alterations in cell numbers in blood of young children and reduced immunological memory in children, while increased inflammation, immunosenescence and reduced naïve memory pools have been described with advanced age (5, 7–10). Interestingly, mouse models have suggested that the mucosal immune system might age more rapidly than the systemic compartment (11).

*Streptococcus pneumoniae* (Spn) is the most common bacterial cause of pneumonia, but usually colonizes the nasopharynx in absence of disease, with colonization frequency decreasing with age (12, 13). Thus, a seeming paradox exists in individuals with advanced age who are at increased risk of pneumococcal disease, but are rarely colonized. On the other hand, children are frequently colonized and infected and are thus the main reservoir for pneumococcal transmission. Mathematical modelling has suggested that the gradual development of adaptive immunity leads to reduced colonization rates with advanced age (14). Mouse models have suggested that Spn colonization is controlled by Th17 cells at the nasal mucosa (15, 16). Using an experimental human pneumococcal challenge model, we recently demonstrated that Spn colonization control in healthy young adults was associated with responses of nasal-resident granulocytes and monocyte recruitment (17, 18).

Here, we aimed to investigate whether immune cell composition at the nasal mucosa are altered in young children and older adults compared to young adults, which could underlie there differential susceptibility to respiratory tract infections. Therefore, we collected minimally-invasive nasal microbiopsies from 207 individuals between 1 to 80 years old and immunophenotyped immune cells using flow cytometry (19). We also investigated circulating innate immune cell function and phenotype and correlated these with mucosal findings. Finally, we investigated the effect of Spn colonization on nasal inflammation by collecting nasal lining fluid and its effect on immune cell composition and activation (20).

## Results

### Composition and activation of nasal immune cells changes with age

Here, we phenotyped nasal cells collected using minimally-invasive nasal microbiopsies (Figure 1). Individuals were grouped as children (1-5 years old, n=43), young adults (18-49 years old, n=121) or older adults (50-80 years old, n=43, Table 1). Samples from children were obtained during planned procedures under general anaesthesia such as dental extractions, while adults were awake for sample collection.

**Table 1.** Cohort characteristics.

|  | Children | Young adults | Older adults |
| --- | --- | --- | --- |
| **Nasal phenotyping sample size** | 43 | 121 | 43 |
| **Age in median years (range)** | 3 (1-5) | 20 (18-49) | 63 (50-80) |
| **Number of females (%)** | 18 (41.2%) | 64 (52.9%) | 26 (60.5) |
| **Number of Spn colonized (%)** | 19 (45.2%*) | 9 (7.4%) | 2 (4.7%) |

\*Streptococcus pneumoniae (Spn) colonization was assessed by classical microbiology.
For one child no nasopharyngeal swab was collected and colonization was not assessed.

**Figure 1.**
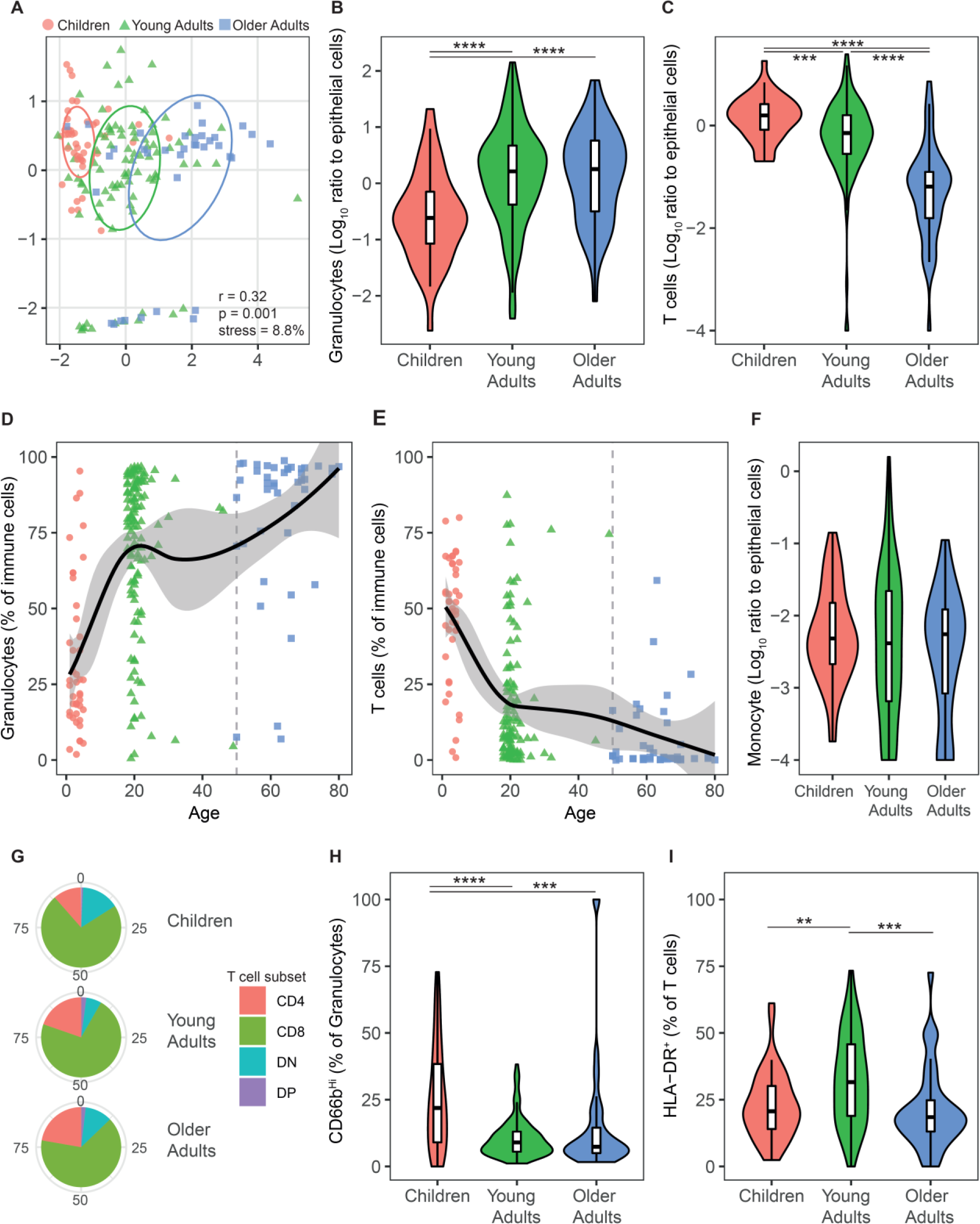
Nasal immune cell populations change drastically with age. A) Multi-dimensional scaling plot based on Euclidian distance, considering immune composition (granulocytes, monocytes and CD4^+^ T, CD8^+^ T and double-negative (DN) T cells as percentage of immune cells) and activation (HLA-DR^+^ and CD66b^Hi^ as percentage of T cells and neutrophils, respectively). Individual children (red circles, n=42), young adults (green triangles, n=86) and older adults (blue squares, n=36) are shown along with 50% confidence intervals. R and p values represent analysis of similarity results and stress depicts Kruskal stress. Violin plots with boxplots showing levels of B) granulocytes and C) T cells normalized to epithelial cells for children (n=43), young adults (n=118) and older adults (n=46). ****p=1.4 x 10^−6^ and p=7.2 x 10^−5^ by Mann-Whitney test comparing granulocytes in children with young adults and older adults, respectively. ***p=0.0006 by Mann-Whitney test comparing T cell levels between children and young adults. ****p=4.5 x 10^−14^ and p=2.4 x 10^−11^ comparing T cell levels in older adults with those in children and with young adults, respectively. Scatterplots showing percentage of D) granulocytes and E) T cells among nasal immune cells. Individual subjects are depicted, a grey vertical dashed line shows the cut-off between young and older adults at age 50 and a locally estimated scatterplot smoothing (loess) curve with 95% confidence interval is plotted. F) Violin plots with boxplots showing levels of monocytes normalized to epithelial cells. G) Pie charts showing the mean levels of T cell subsets (CD4^+^, CD8^+^, double-negative (DN) and double-positive (DP) per age group (children, n=43; young adults, n=109; older adults, n=42). H) Violin plots with boxplots showing levels of CD66b^Hi^ granulocytes for children (n=43), young adults (n=121) and older adults (n=41). ****p=2.3 x 10^−6^ and ***p=0.0004 by Mann-Whitney test comparing children with young adults and older adults, respectively. I) Violin plots with boxplots showing levels of HLA-DR^+^ T cells for children (n=43), young adults (n=109) and older adults (n=38). **p=0.001 and ***p=0.0002 comparing young adults with children and older adults by Mann-Whitney test, respectively.

Nasal cell populations exhibited a significant shift with age (Figure 1A). The capacity to maximally discriminate groups occurred at an age cut-off between young and older adults at 50 years (see Figure S1 in the SI Appendix). Levels of granulocytes were 6.7x and 7.4x lower in children than in young adults and older adults, respectively (Figure 1B). Conversely, nasal T cell levels were 16.1x and 8.1x lower in older adults than in children or young adults, respectively (Figure 1C). As a proportion of nasal immune cells, neutrophils increased, and T cells decreased with age (Figure 1D, E). Monocytes were rare at the nasal mucosa and did not change in frequency with age (Figure 1F). Among T cells, CD8^+^ T cells were the most abundant subset in all age groups (Figure 1G). CD4^−^ CD8^−^ T cell numbers and CD8^+^ T cell numbers were 4.8x and 2.0x increased in children compared to young adults, respectively (see Figure S2 in the SI Appendix). CD4^+^ T cell numbers were similar between children and young adults and CD4^+^ T cells thus increased in frequency among T cells in adults (Figure 1G and see Figure S2 in the SI Appendix). In children, granulocytes were fewer, but exhibited higher expression of CD66b, a marker of degranulation, than in adults (Figure 1H) (21). T cell activation status was also affected by age, with young adults showing increased levels of human leukocyte antigen – DR isotype (HLA-DR)^+^ T cells compared to children and older adults (Figure 1I).

### Circulating monocytes from young children have reduced CCL2 production upon *Streptococcus pneumoniae* stimulation

We then investigated how innate immune function was affected by age. As the numbers of nasal cells collected using curettes precluded the conduction of functional assays, blood was used. To measure monocyte function, we stimulated whole blood from 43 individuals for 4 hours with heat-killed Spn and assessed production of C-C motif chemokine ligand 2 (CCL2), interleukin-10 (IL-10), IL-6 and tumor necrosis factor alpha (TNF) (see Figure S3 in the SI Appendix). Monocytes from children displayed impaired production of CCL2 upon stimulation compared to adults, while production of IL-6 and TNF was similar (Figure 2A). Little or no of the anti-inflammatory cytokine IL-10 was induced, demonstrating a pro-inflammatory response of blood monocytes upon Spn stimulation.

**Figure 2.**
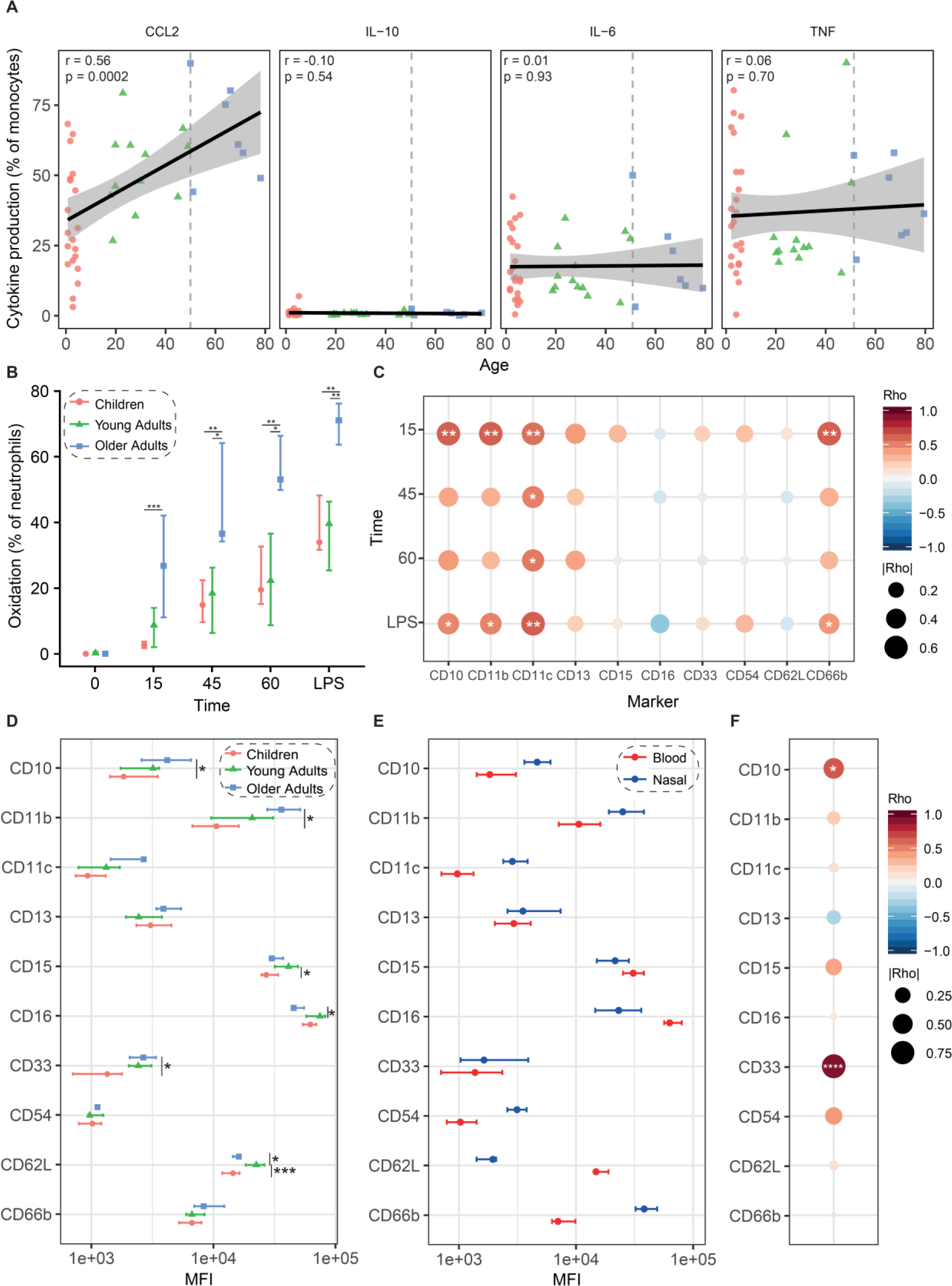
Changes in innate immune function with age. A) Scatterplots showing percentage of monocytes that are producing CCL2, IL-10, IL-6 or TNF following a two-hour stimulation with heat-killed Streptococcus pneumoniae. Individual children (red circles, n=23), young adults (green triangles, n=13) and older adults (blue squares, n=7) are depicted. Black line and shaded area depict linear regression fit and 95% confidence intervals. R and p represent rho and p-value from Pearson’s correlation test, respectively. B) Neutrophil oxidation capacity was assessed using a phagocytic bead assay at 4 timepoints, including a positive control where LPS was added for 45 minutes. Median and interquartile range are shown for children (red circles, n=11), young adults (green triangles, n=6) and older adults (blue squares, n=5). *p<0.05, **p<0.01, ***p<0.001 by Mann-Whitney test comparing two groups. C) Correlation matrix for oxidative capacity and log-transformed blood neutrophil surface marker expression using Pearson correlation test (n=21). Circle colour and size represent rho and absolute rho value, respectively. *p<0.05, **p<0.01 by Pearson correlation test D) Mean fluorescent intensity (MFI) of surface markers on blood neutrophils per age group. Median and interquartile range are shown for children (red circles, n=13), young adults (green triangles, n=8) and older adults (blue squares, n=5). *p<0.05, ***p<0.001 by Mann-Whitney test. E) Comparison of marker expression on neutrophils from blood (red) and nose (blue) for paired volunteers (n=11 children and 2 young adults). F) Correlation matrix for log-transformed marker expression on blood and nasal neutrophils using Pearson correlation test (n=13). Circle colour and size represent rho and absolute rho value, respectively. *p=0.018, ****p=1.1×10^−5^ by Pearson correlation test.

### Blood neutrophils change functionally and phenotypically with age

Neutrophil phagocytic and oxidative capacities were measured using a whole blood reporter bead assay (see Figure S4 in the SI Appendix) (22). In parallel, neutrophils were immunophenotyped using a panel of ten maturation and activation markers (see Figure S4 in the SI Appendix). Neutrophils of older adults displayed increased uptake and oxidation, while no significant differences were present between neutrophils of children and young adults (Figure 2B). The oxidative capacity of neutrophils positively correlated with expression of the activation and maturation markers CD10, CD11b, CD11c and CD66b (Figure 2C). In addition, neutrophil surface levels of CD10, CD11b and CD33, but not CD11c and CD66b, were significantly increased in older adults compared to children (Figure 2D). Blood neutrophils in young adults had increased expression of CD62L compared to children and older adults and increased expression of CD15 and CD16 compared to children and older adults, respectively. To compare blood and mucosal neutrophils, paired neutrophils were phenotyped for 13 individuals (Figure 2E). Nasal neutrophils had increased surface expression of CD10, CD11b, CD11c, CD54 and CD66b, while CD62L and CD16 expression were lost. Neutrophils at the nasal mucosa thus exhibit an activated phenotype as previously described for neutrophils in bronchoalveolar fluid (23). Of all markers, only CD10 (r=0.64) and CD33 (r=0.92) positively correlated between the two compartments, indicating that blood neutrophil phenotype does not accurately reflect mucosal neutrophil phenotype on an individual level (Figure 2F).

### *Streptococcus pneumoniae* colonization causes inflammation in children

Finally, we investigated how colonization with Spn affects nasal immune populations and responses by measuring levels of thirty cytokines and chemokines in nasal lining fluid collected from adults (n=37, none colonized by Spn) or children (n=49, 22 of whom colonized by Spn). Cytokine levels were similar between adults and children not colonized with Spn, but different in children colonized with Spn (Figure 3A). Levels of granulocyte-colony stimulating factor (G-CSF), IL-15, IL-6, CCL3, basic fibroblast growth factor (FGF-basic), IL-17A, IL-10, CCL5, TNF and IL-5 were significantly elevated in colonized children compared to adults (Figure 3B). Among these, CCL3 and IL-6 showed a positive association with pneumococcal load (Figure 3C). G-CSF was the only protein also increased in non-colonized children compared to adults. Interleukin-1 receptor antagonist (IL-1RA) and IL-13 were decreased in both colonized and non-colonized children compared to adults, while IL-7 was lower in only non-colonized children compared to adults. There were no significant differences between colonized and non-colonized children, but these groups showed a relatively large inter-individual variation compared to adults. This is in accordance with previous findings that the gut microbiome shows greater variation in young children than in adults (24). Subsequently, we assessed neutrophil degranulation by measuring nasal levels of myeloperoxidase (MPO), which were increased in both non-colonized (1.5x, p = 0.02) and colonized (4.6x, p = 0.0005) children compared to adults (Figure 3D) (25). However, levels of MPO were not significantly affected by Spn colonization status or load in children (Figure 3D, E). Nonetheless, the increased levels of MPO in children compared to adults confirms the increased expression of CD66b on nasal granulocytes in children. Indeed, MPO levels correlated on an individual level with granulocyte activation but not granulocyte numbers (Figure 3F, G).

**Figure 3.**
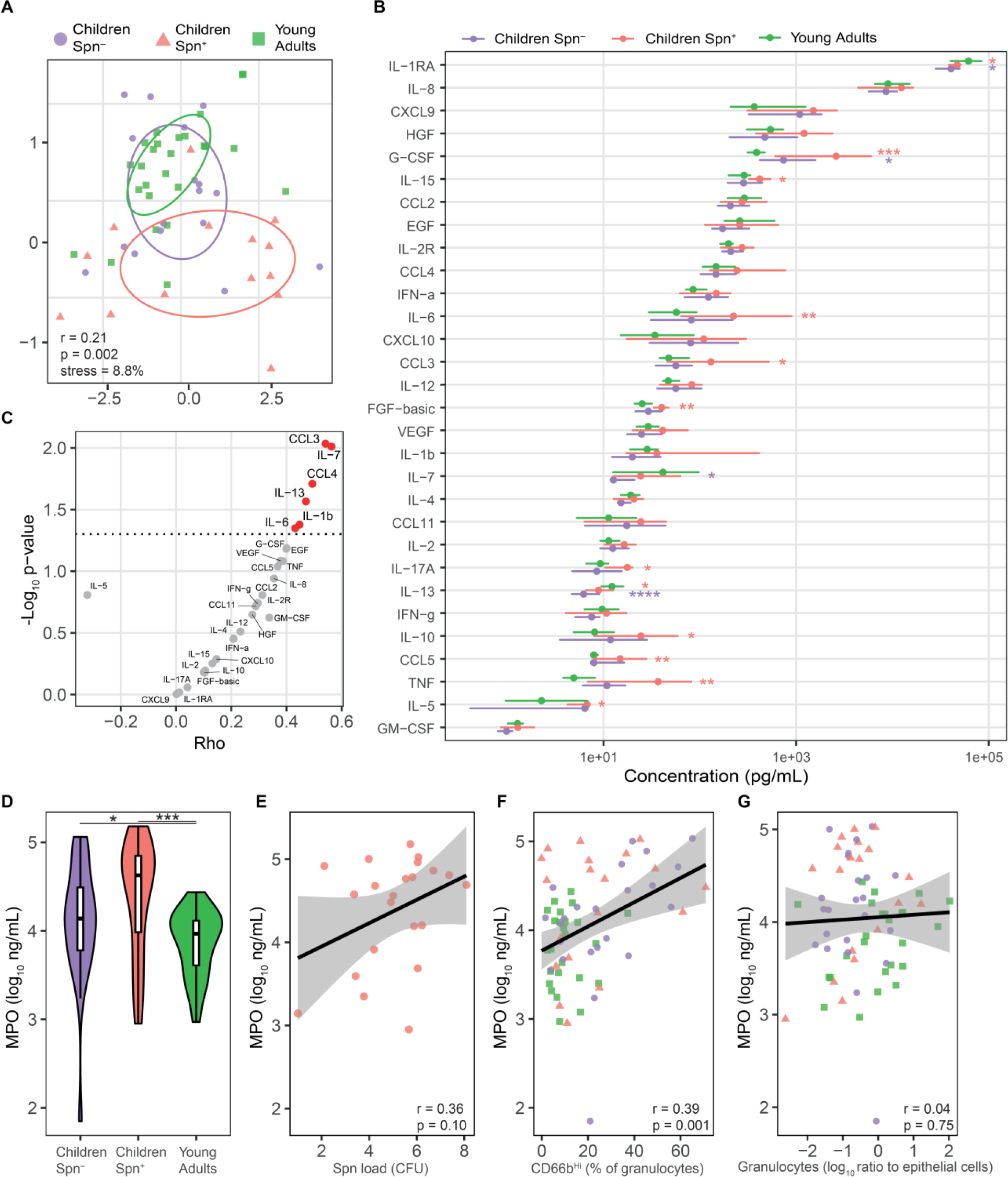
Nasal cytokine responses to pneumococcal colonization. A) Multi-dimensional scaling plot based on Euclidian distance, considering log-transformed concentrations of 30 nasal cytokines. Individual Spn not-colonized children (Spn^−^ purple circles, n=17), Spn colonized children (Spn^+^ red triangles, n=14) and young adults (green squares, n=26) are shown along 50% confidence intervals. R and p values represent analysis of similarity results and stress depicts Kruskal stress. B) Median and interquartile range of the concentrations for each of the 30 cytokines are shown for not-colonized children (purple, n=27), colonized children (red, n=22) and young adults (n=37). *p<0.05, **p<0.01, ***p<0.001, ****p<0.0001 by Mann-Whitney test compared to young adults, followed by Benjamini-Hochberg correction for multiple testing, with colour indicating significantly altered group. C) Volcano plot showing Pearson correlation results between log-transformed pneumococcal load for colonized children with log-transformed cytokine concentration, not corrected for multiple testing. Significantly correlating cytokines are depicted in red. D) Violin plots with boxplots showing concentration of myeloperoxidase (MPO) for not-colonized children (purple, n=27), colonized children (red, n=22) and young adults (green, n=36). *p=0.022, ***p=0.0005 by Mann-Whitney test. Scatterplots show correlation between log-transformed concentration of nasal MPO with E) log-transformed pneumococcal load, F) percentage of CD66b^Hi^ granulocytes and g) log-transformed levels of nasal granulocytes (normalized to epithelial cells). Individuals are shown, and line and shaded area represent linear regression and 95% confidence interval, respectively. R (Pearson rho) and p (p-value) are shown for each correlation.

### *Streptococcus pneumoniae* colonization is not associated with altered nasal immune cells in children

Despite the increased cytokine production during Spn colonization, no clear differences in immune cell levels were apparent between colonized and non-colonized children (Figure 4A-C). Monocytes, which are recruited to the nose of adults experimentally colonized with pneumococcus, were not affected by Spn colonization (Figure 4D) (17). While Spn colonization was not associated with neutrophil activation levels, T cell activation was increased in Spn-colonized children compared to non-colonized children (29.3% versus 18.1% of HLA-DR^+^ T cells, Figure 4E, F).

**Figure 4.**
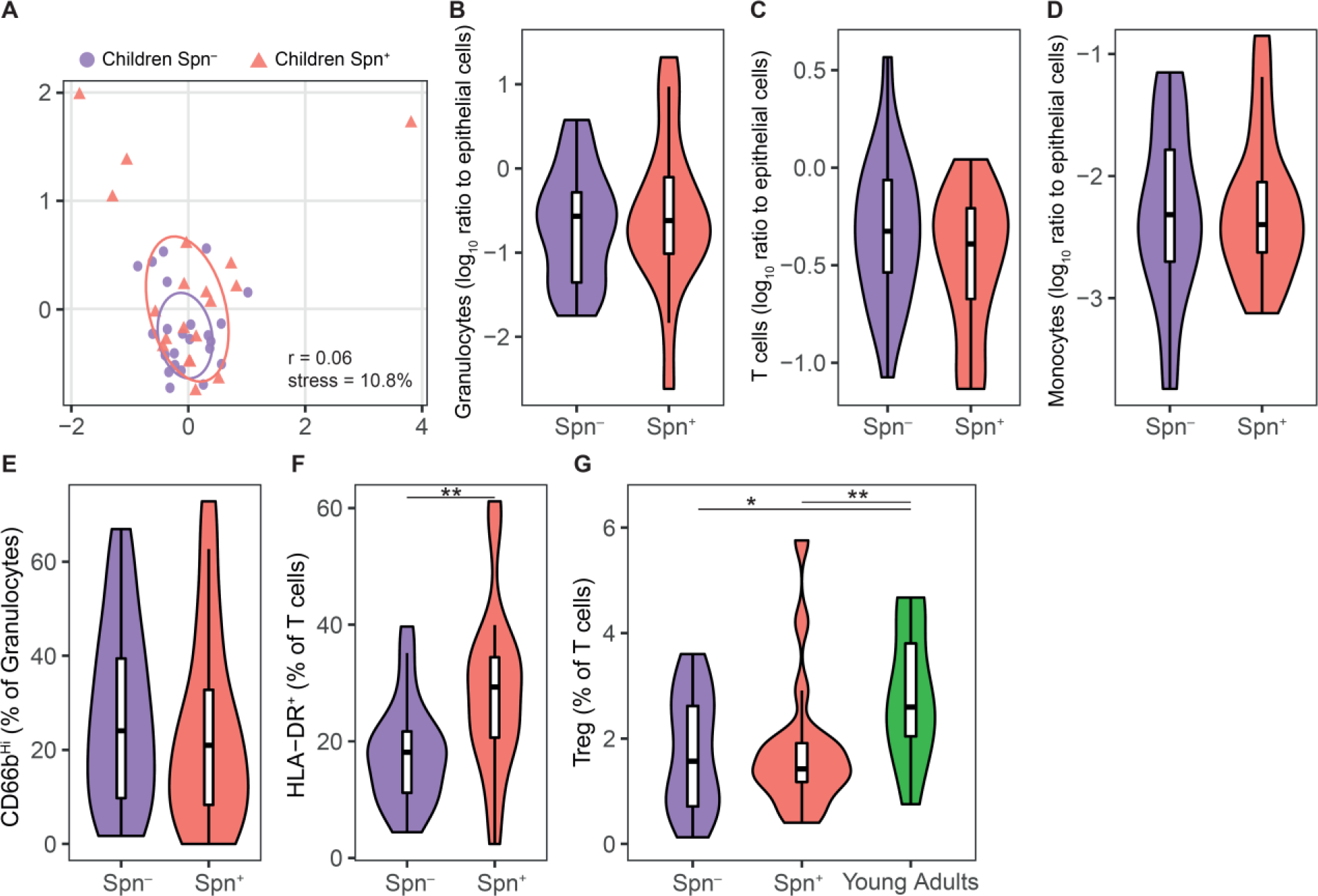
Nasal immune cell populations are not affected by Spn colonization in children. A) Multi-dimensional scaling plot based on Euclidian distance, considering immune composition (granulocytes, monocytes and CD4^+^ T, CD8^+^ T and double-negative (DN) T cells as percentage of immune cells) and activation (HLA-DR^+^ and CD66b^Hi^ as percentage of T cells and neutrophils, respectively). Individual non-colonized children (Spn^−^ purple circles, n=23) and colonized children (Spn^+^, red triangles, n=19) are shown along with 50% confidence intervals. R represents the analysis of similarity results and stress depicts Kruskal stress. Violin plots with boxplots showing numbers of B) granulocytes, C) T cells or D) monocytes normalized to epithelial cells for non-colonized children (Spn^−^, purple, n=23) and colonized children (Spn^+^, red, n=19). Violin plots with boxplots showing levels of E) CD66b^Hi^ granulocytes or F) HLA-DR^+^ T cells. **p=0.005 by Mann-Whitney test. G) Violin plots with boxplots showing numbers of regulatory T cells (Tregs) for non-colonized children (Spn^−^, purple, n=23), colonized children (Spn^+^, red, n=19) and non-colonized young adults (green, n=11). *p=0.036, **p=0.0098 comparing young adults with colonized and non-colonized children by Mann-Whitney test, respectively.

## Discussion

Here, we investigated how the composition of immune cells at the nasal mucosa is altered with age, with only young adults having an immune profile with an abundance of both granulocytes and T cells. Granulocytes were depleted in children, while there was a paucity of T cells in older adults.

As neutrophils have a critical concentration threshold for effective bacterial killing, it is possible the reduced number of nasal granulocytes is associated with the increased susceptibility of children to respiratory tract colonization and infections (26). The reduced expression of the adhesion molecules CD62L, CD11b and CD15, which are important for extravasation and trafficking to tissues (27), on blood neutrophils from children could explain their limited numbers at the nasal mucosa. Nasal IL-8 levels, which is important for neutrophil migration, were not different between adults and children. We did not, however, investigate expression of chemokine receptors as CXCR1 and 2, the receptors for IL-8, on circulating neutrophils and it cannot be excluded that differences in surface expression of these markers exist between children and adults (28).

Despite having reduced numbers of nasal granulocytes, the granulocytes in children were activated to a higher degree than in young adults, as shown by increased levels of CD66b expression and increased levels of MPO. Possible explanations for this include a different effect of migration into tissues, which can activate neutrophils *per se*, and altered activation by microbiota in children compared to adults. Indeed, surface marker expression between blood and nasal neutrophils correlated poorly on an individual level, with the exception of CD10 and CD33. This highlights that the study of immune cells in the circulation potentially does not reflect the same cell type at mucosal sites.

Circulating neutrophil responses using the bead reporter assay were increased in older adults compared to younger adults, in agreement with previous reports on neutrophils from elderly individuals (29). Our observations also corroborate findings from mouse models where aged neutrophils show an increased phagocytic capacity compared to non-aged neutrophils, which was associated with elevated expression of CD11b (27). In contrast, previous studies have shown that the antibody-mediated opsonophagocytic capacity of neutrophils is reduced in the elderly with reduced observed responses to *Staphylococcus aureus*, but not *Escherichia coli* (30, 31).

We observed that T cell levels were reduced in older adults, which was already apparent around the age of 50, before the increased susceptibility to infections becomes apparent. As tissue-resident memory T cells are crucial for protection against infections at mucosal surfaces, this lack of mucosal T cells could provide an explanation for the increased susceptibility to respiratory infections in the elderly (32). Although vaccine efficacy drops with advanced age (33), the development of vaccines that increase tissue-resident memory T cells might thus be particularly beneficial for the elderly. In that light it would be interesting to further characterize these T cells at the mucosa to investigate which specific cells are lost, although this would require access to large tissue samples such as biopsies.

Spn colonization led to increased nasal inflammation in children, in contrast to what we previously observed in experimentally colonized adults (17). This could explain the increased transmission potential of children, as inflammation was shown to augment transmission in murine models (34). Although the molecular mechanisms that associate with increased inflammation upon colonization in children remain unclear, it is likely not uniquely due to higher pneumococcal density in children as many cytokines only poorly correlated with Spn density. It is not impossible that the concurrent presence of other bacterial and viral factor was associated inflammation but also predisposing to Spn colonization.

Spn colonization had limited effect on the nasal immune cell composition or activation status in children, with the exception of HLA-DR expression on T cells. This corroborates the observed increase in nasal levels of prototypic T cell cytokines as IL-17A, IL-5, IL-10 and CCL-5 in colonized children. Levels of MPO and CD66b expression on neutrophils were not affected by Spn colonization in children. This is in contrast with young adults experimentally inoculated by Spn, who show an increase in MPO levels following colonization (17). Possible explanations for this discrepancy are the reduced total number of granulocytes in children, an increased baseline neutrophil activation in non-colonized children compared to adults, or presence of low numbers of Spn that are not detected by classical microbiology but could still activate neutrophils.

In addition, Spn colonization in children was not associated with increased levels of monocytes, which are important for Spn clearance (17, 35). Indeed, nasal CCL2 levels, which mediates monocyte recruitment (17, 35), were not increased in colonized children. Interestingly, blood monocytes from children showed an impaired production of CCL2 upon Spn stimulation. This was specific as production of TNF and IL-6 was not affected. Previously it was shown that infant mice also have a limited monocyte recruitment following Spn colonization, leading to an inability to clear Spn colonization, although this was associated with microbiota-driven increased baseline CCL2 levels (36).

In contrast to our findings, in one previous study in which nasal aspirates were collected in children with acute otitis media, upper respiratory infections or without infection, recruitment of neutrophils to the nasopharynx correlated with Spn density (37). However, the previous study used qPCR for the genes *CD16*, *CD18* and *CD62L* on nasal aspirate pellets to quantify neutrophils, which might not as accurately measure neutrophil counts as flow cytometry, since qPCR reflects both the cellular composition and gene expression levels of individual cells. Moreover, the previous study found increased neutrophil count especially during otitis media, which also associated with increased Spn density, while we studied immune composition in the absence of infection.

One previously postulated explanation for the increased susceptibility to Spn colonization in children is an increased Treg/Th17 ratio in children compared to adults (38, 39). However, IL-17A levels were elevated in nasal fluid of colonized children (Figure 3B). In addition, levels of nasal Tregs were not affected by colonization state and were lower in children than in young adults (Figure 4G).

A limitation of this cross-sectional observational study was that we focused on three groups: young children, young adults and older adults. Consequently, we have no older children from 6-17 years and we also have few adults between 30-50 years. This makes it hard to detect at which ages nasal immune profiles start shifting.

In conclusion, we observed severe and dynamic alterations in mucosal immunity with age, highlighting the need for measuring mucosal responses in target populations when investigating host-pathogen interactions and vaccine-induced immunity, especially in the young and elderly.

## Methods

### Study design

We recruited individuals between 1–80 years of age in a series of studies (ISRCTN85509051, ISRCTN10948363, ISRCTN16993271, ISRCTN68323432 and ISRCTN76456378). Some of the subjects in this manuscript (the young adults cohort) were originally described previously (40). For a subset of the young adults, cytokines and nasal cell data have been used in another manuscript deposited on a pre-print server (18), although the results do not overlap with those reported here. All adults were healthy and inclusion criteria common to all adult studies were: capacity to give informed consent, aged >18 years and speak fluent English. Children awaiting a procedure requiring general anaesthesia: dental extraction (44%), MRI (36%), orthopaedic surgery (14%) and plastic surgery (6%), were recruited. Inclusion criteria were: aged 1-5 years, capacity of a parent of the participant to give informed consent and speak fluent English. Exclusion criteria and sample collection details for adults and children are provided in an online data supplement.

### Ethics statement

All adult volunteers and a parent of children involved in the study gave written informed consent and research was conducted in compliance with all relevant ethical regulations. Ethical approval was obtained from the East Liverpool NHS Research Ethics Committee, reference numbers: 17/NW/0663, 16/NW/0031, 17/NW/0029, 15/NW/0931 and 14/NW/1460.

### Flow cytometry analysis

All flow cytometry samples were acquired on a LSRII flow cytometer (BD) and analysed using Flowjo X (Treestar). Compensation matrices were set using compensation beads (BD Biosciences) and ArC™ Amine Reactive Compensation beads (Thermofisher) and manually inspected for representative samples. All antibodies were titrated and fluorescence minus one controls were used to verify specificity of signal. Additional detail on immunophenotyping of nasal cells, neutrophil phenotyping and monocyte and neutrophil functional assays is provided in an online data supplement.

### Luminex analysis of nasal lining fluid

Cytokines were eluted from stored Nasosorption™ filters using 100µL of Luminex assay buffer (ThermoFisher) by centrifugation, then the eluate was cleared by further centrifugation at 16,000 x G, as described previously (17, 19). Concentrations of 30 cytokines were measured using the 30-plex magnetic human Luminex cytokine kit (all using lot ID 1805187A, ThermoFisher). Samples were measured on a LX200 (Luminex) and analysed with xPonent3.1 software (Luminex) following manufacturer’s instructions. Samples were analysed in duplicates and analytes with a CV > 50% were excluded.

### Myeloperoxidase (MPO) ELISA of nasal lining fluid

Levels of myeloperoxidase were determined using the Human Myeloperoxidase DuoSet ELISA Kit (R&D Systems) as per manufacturer’s instructions. Plates were read on a FLUOstar^®^ Omega machine (BMG Labtech) and data was analysed with Mars data analysis software version 3.1 following manufacturer’s instructions. Samples were analysed in duplicates and samples with a CV > 20% were excluded.

### Statistical analysis

Statistical analyses were performed using R software (version 3.5.1). Two-tailed statistical tests were used throughout the study. Mann-Whitney tests were used to compare groups and multiple correction testing (Benjamin-Hochberg) was applied for Luminex analysis. Correlations were assessed using Pearson’s correlation test using either raw or log-transformed values. Differences were considered significant at p < 0.05.

## Abbreviations

Spn: *Streptococcus pneumonia*
MPO: Myeloperoxidase
LPS: Lipopolysaccharide

## Acknowledgements

This work was supported by Bill and Melinda Gates Foundation (OPP1117728 awarded to Daniela Ferreira), the Medical Research Council (MR/K01188X/1 awarded to S.G Gordon) and we acknowledge the support of the National Institute for Health Research Clinical Research Network. This work was also supported by the Liverpool School of Tropical Medicine Director Catalyst Fund which was funded by the Wellcome Trust Institutional Strategic Support Fund 3 (204806/Z/16/Z) and the Liverpool School of Tropical Medicine Internal Funding (awarded to Simon Jochems). Flow cytometry acquisition was performed on a BD LSR II cytometer and BD FacsAria III Cell Sorter funded by a Wellcome Trust Multi-User Equipment Grant (104936/Z/14/Z).

We would like to thank all volunteers for participating in this study and R. Robinson, C. Lowe, L. Lazarova, K. Piddock and I. Wheeler for clinical support. Unencapsulated *Streptococcus pneumoniae* was a gift by J. Brown. Daniela Ferreira and Simon Jochems are members of the Human Infection Challenge Network for Vaccine Development (HIC-Vac), which is funded by the Global Challenges Research Fund Networks in Vaccines Research and Development, which was co-funded by the Medical Research Council and Biotechnology and Biological Sciences Research Council.

