## Supplemental Data for "Dynamic changes in innate immune and T cell function and composition at the nasal mucosa across the human lifespan"

### Supplemental Methods

#### Exclusion criteria

Exclusion criteria common to all adult studies included: current involvement in another study unless observational or in follow-up phase (non-interventional), close contact with children under 5 years, current regular smoker, >10 pack years smoking history, asthma or respiratory disease, pregnant, on medication that may affect the immune system in any way e.g. steroids, steroid nasal spray, regularly taking acetylsalicylic acid (aspirin), current acute severe febrile illness and taking long term antibiotics.

Exclusion criteria for the paediatric study were: taking daily medications that may affect the immune system, having received antibiotics in the preceding 28 days, history of respiratory infections requiring hospitalization, involved in another clinical trial unless observational or in follow-up (non-interventional) phase, having a disease or syndrome associated with altered immunity, undergoing procedure related to infection, asthma and current severe acute respiratory infection. Although not an exclusion criterion, none of the children had ever had a tonsillectomy.

#### Sample collection

In adults, all samples were collected while awake, while samples from children were collected after onset of general anaesthesia but prior to start of their planned procedure. Nasal microbiopsies (ASL Rhino-Pro©, Arlington Scientific) and nasal lining fluid (Nasosorption™, Hunt Developments) samples were collected and stored at -80°C as previously described (1, 2). For adults, a nasal wash was performed to assess pneumococcal colonization by classical microbiology as described (3). For children, a nasopharyngeal swab was collected and stored in 1mL of skim milk, tryptone, glucose, and glycerine (STGG) medium. Pneumococcal colonization was assessed using classical microbiology as done for adults. Whole blood was collected in lithium heparin tubes, up to 3mL for children and up to 36mL for adults. Blood was kept under agitation at room temperature and used within 2 hours of collection.

**Immunophenotyping of nasal cells**

Nasal cells were phenotyped using flow cytometry as previously described (1, 4). In brief, cells were dislodged from microbiopsies and stained for 15 minutes on ice with LIVE/DEAD® Fixable Violet Dead Cell Stain (ThermoFisher) and then stained for a further 15 minutes with an antibody cocktail composed of the following antibodies (antibody clones in parentheses): Epcam-PE (9C4), HLADR-PECy7 (L243), CD16-BV711 (3G8), CD66b-FITC (G10F5), CD3-APCCy7 (SK7), CD45-BV510 (HI30), CD4-BV605 (RPA-T4), CD8-BV786 (SK1, all from Biolegend) and CD14-PerCPCy5.5 (MφP9, BD Biosciences). In addition, to subsets of samples we added additional markers CD25-PE/Dazzle (M-A251), GARP-APC (LRRC32), TCRVα7.2-PE/Dazzle594 (3C10), CD107a-BV650 (H4A3), CD279-BV711 (EH12.2H7), TCRVα7.2-BV711 (3C10), CD1c-PE/Dazzle594 (L161, all from Biolegend), BDCA2-APC (AC144, Miltenyi Biotec) and BDCA3-BV711 (1A4, BD Biosciences) in different combinations. At the end of the incubation, cells were washed, filtered over a 70μm filter (ThermoFisher) and spun down (440 x G for 5 min). Then, cells were resuspended in 200μL of Cell Fix (BD Biosciences) and acquired on a flow cytometer (LSRII, BD). Samples with less than 500 immune cells or 250 epithelial cells were excluded from further analysis. A total of 67 samples were thus excluded

(7/50,14%); samples from children, (39/160, 24%); samples from young adults and (21/64, 32%); samples from older adults)

**Whole blood and nasal neutrophil immunophenotyping**

Nasal cells and blood samples were stained using LIVE/DEAD® Fixable Violet Dead Cell Stain (ThermoFisher, L34964) for 15 minutes and then for an additional 15 minutes with an antibody cocktail containing the following antibodies (antibody clones in parentheses): CD66b-FITC (G10F5), CD33-PE (P67.6), CD54-PE/Dazzle (HA58), CD11c-PerCPCy5.5 (Bu15), CD11b-PECy7 (ICRF44), CD10-APC (HI10a), CD45-BV510 (HI30), CD15-BV605 (W6D3), CD62L-BV650 (DREG-56), CD16-BV786 (3G8), CD13-APCCy7 (WM15) and Epcam-PE (9C4, all from Biolegend). The epithelial marker Epcam was included in the staining of nasal cells but not in the staining of blood. Whole blood was stained at room temperature, while nasal cells were stained on ice. At the end of the incubation nasal cells were washed and filtered over a 70µm filter (ThermoFisher). Whole blood was lysed using FACS™ Lysing Solution (BD Biosciences) according to manufacturer's instructions at the end of the incubation period. Finally, cells were spun down (440 x G for 5 min), resuspended in 200µL of Cell Fix (BD Biosciences) and acquired on a flow cytometer (LSRII, BD).

**Monocyte cytokine production after Spn6B stimulation**

Monocyte cytokine production from whole blood was assessed similar to previously described (5). Fresh whole blood samples (200µL) were stimulated in presence or absence of heat-killed, unencapsulated *Streptococcus pneumoniae* at 5µg/mL in a 96 well-plate with rounded bottom for 2 hours at 37°C and 5% $CO_2$. Cytokines were retained within the cells by the addition of GolgiPlug (BD Biosciences) followed by stimulation for 2 more hours. Post incubation, samples were detached from the wells by adding 2mM EDTA and incubating for 10 minutes at room temperature. Samples were then lysed and fixed using FACS™ Lysing Solution (BD Biosciences) for 10 minutes at room temperature. Samples were then spun down, resuspended in CTL-Cryo™ ABC (Fisher Scientific) and stored at -80°C until batch staining and acquisition. Cells were thawed on ice, washed with 3mL PBS and permeabilized by a 10-minute incubation with Perm/Wash™ (BD Biosciences) on ice. Subsequently, samples were spun down and stained for 45 min protected from light at 4°C with an antibody cocktail containing CD66b-FITC (G10F5), MCP1-PE (2H5), IL6-APC (MQ2-13A5), CD450BV510 (HI30), TNF-α-BV605 (MAb11), IL10-BV786 (JES3-9D7, all from Biolegend) and CD14-PerCPCy5.5 (MφP9, BD Biosciences). Samples were washed and resuspended in 300uL PBS and acquired on a LSRII flow cytometer (BD).

**Neutrophil phagocytosis assay**

Neutrophil function was assessed in fresh whole blood samples using labelled beads as described previously (6). In short, carboxylated silica beads (PSI-3.0COOH, Kisker Biotech) were labelled with a reporter fluorochrome 2’,7’-dichloro-dihydro-fluorescein diacetate (Oxyburst Green, D-2935) and a calibrator fluorochrome (Alexa Fluor 405SE, A30000). A volume of 100µL of whole blood was mixed with labelled beads and incubated for 0, 15, 45 and 60 minutes protected from light at 37°C on an orbital shaker. LPS (100ng/mL, Sigma Aldrich) was added as a positive control for 45 minutes. Samples were then placed on ice to stop bead phagocytosis and oxidation and samples were stained using LIVE/DEAD® Fixable Far Read Dead Cell Stain (ThermoFisher, L34973) and an antibody cocktail composed of CD66b-PerCPCy5.5 (G10F5) and CD16-PECy7 (3G8, All from Biolegend). Then, samples were lysed using FACS™ Lysing Solution (BD Biosciences), washed and resuspended in 300uL PBS. A total of 30,000 neutrophils were acquired from each sample.

### Supplemental Figures

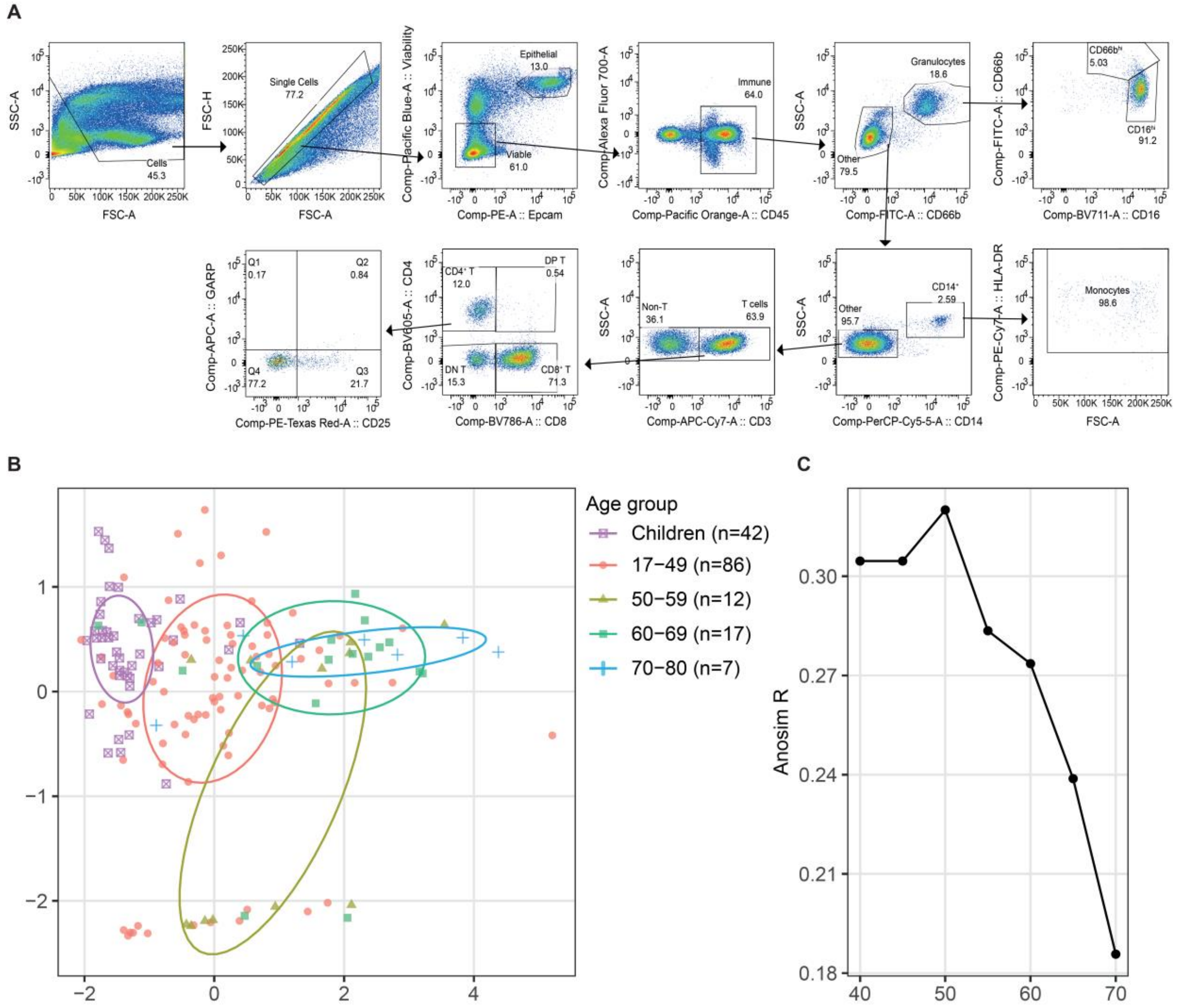


**Figure S1. Nasal cell phenotyping.** A). The applied flow cytometric gating strategy is shown for one representative paediatric sample. B) Multi-dimensional scaling plot based on Euclidian distance, considering immune composition (granulocytes, monocytes and $CD4^+$ T, $CD8^+$ T and double-negative (DN) T cells as percentage of immune cells) and activation (HLA-$DR^+$ and $CD66b^{Hi}$ as percentage of T cells and neutrophils, respectively). Individual children (purple crosses), 17-49-year old's (red circles), 50-59-year old's (yellow triangles), 60-69-year old's (green squares) and 70-80-year old's (blue crosses) are shown along with 50% confidence intervals. C) Anosim R statistic is shown for an incrementally increasing cutoff between young and old adults from 40 to 70 years, peaking at age 50.

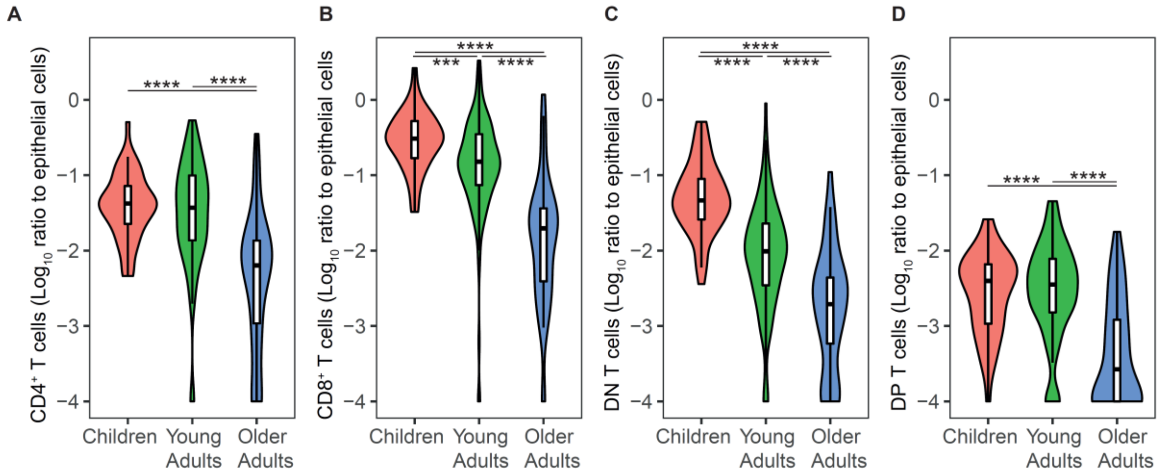


**Figure S2. Nasal T cell subsets change with age.** Violin plots with boxplots showing levels of A) $CD4^{+}$, B) $CD8^{+}$, C) double-negative (DN) and D) double-positive (DP) T cells normalized to epithelial cells for children (n=43, red), young adults (n=109, green) and older adults (n=45, blue). ***p<0.001 and ****p<0.0001 by Mann-Whitney test comparing levels between groups.

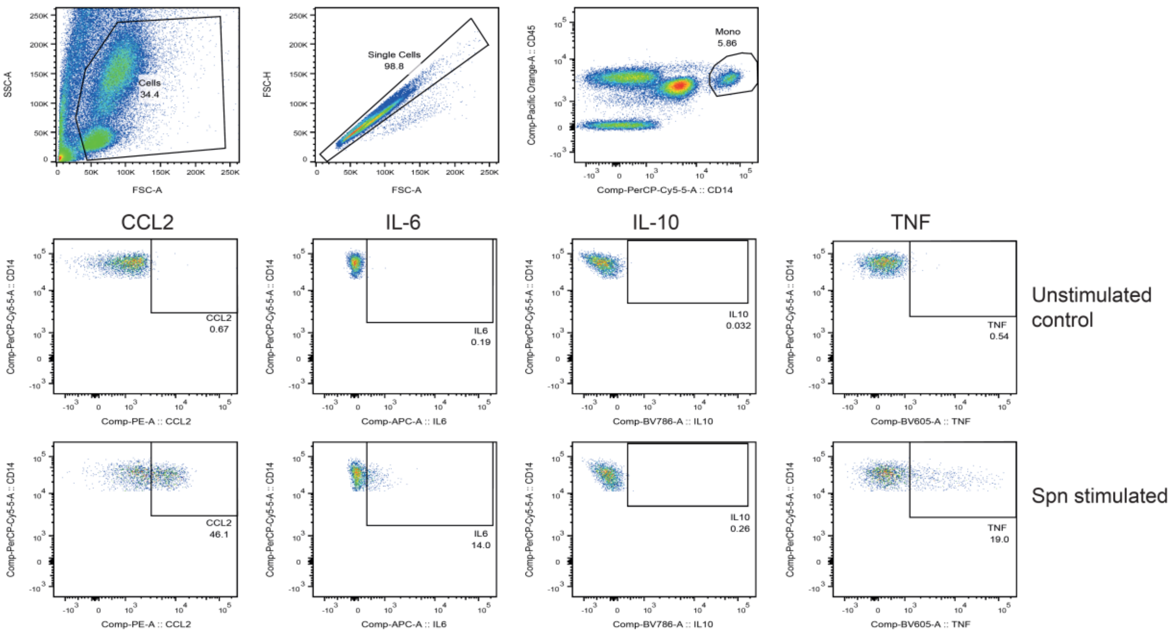


**Figure S3. Representative results for monocyte immune function assay.** Intracellular cytokine staining in monocytes from unstimulated or pneumococcus-stimulated whole blood.

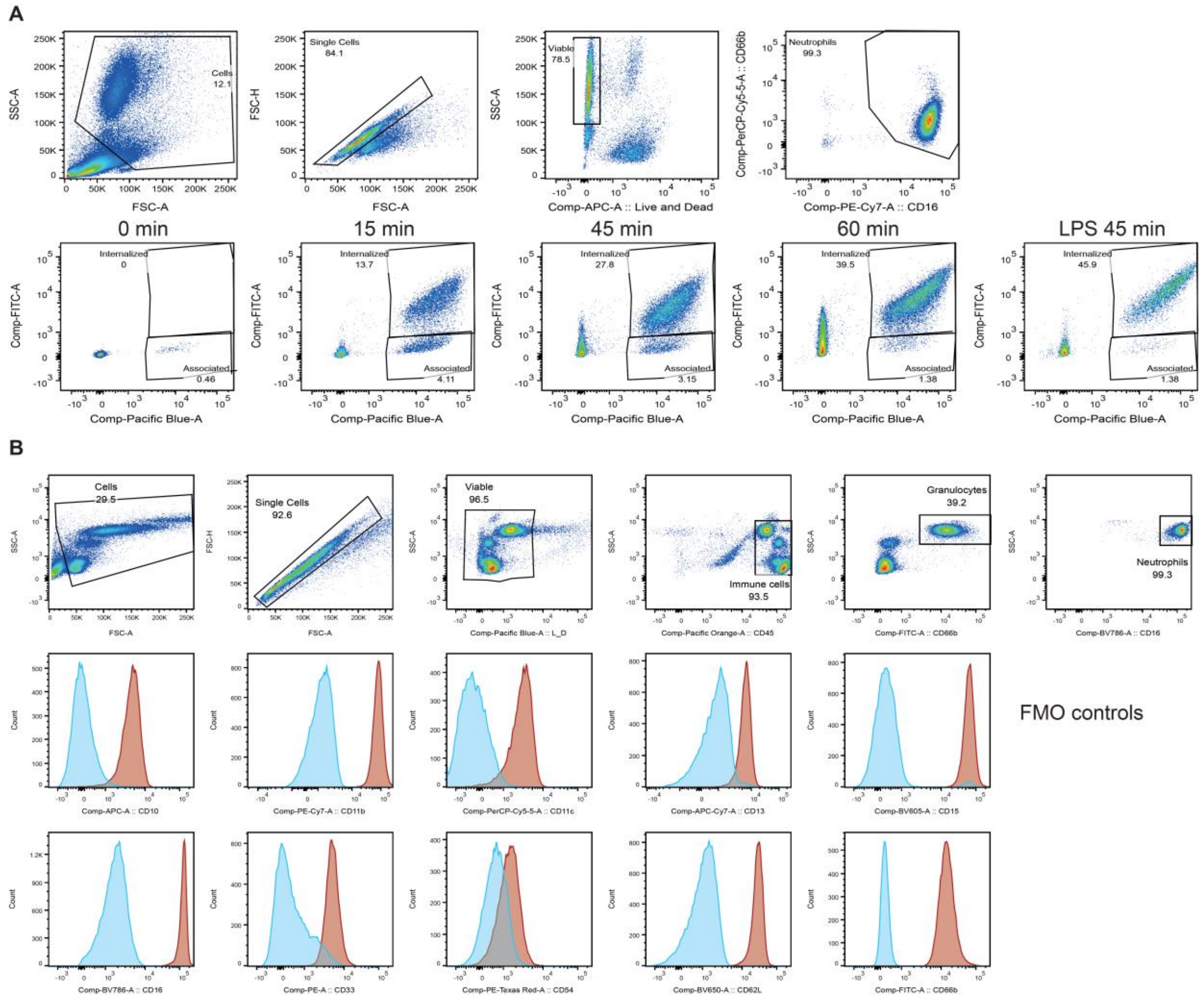


**Figure S4. Representative results for neutrophil immune function assay.** A) Gating strategy for the neutrophil bead reporter assay. Flow cytometry data analysis is shown for one representative paediatric sample. Neutrophil bead association and oxidation at 0, 15, 45, 60 minutes or 45 minutes in presence of lipopolysaccharide (LPS) are shown. B) Neutrophil gating strategy and surface marker expression on whole blood. Red histograms show signal in neutrophils and blue histograms represent fluorescent minus one (FMO) controls for each of the markers.
